# Dual cryo-radiofrequency ablation enhances lesion depth in beating human left ventricular preparations

**DOI:** 10.1101/469882

**Authors:** Matthew S. Sulkin, Jacob I. Laughner, Michael Rogge, Jonathan M. Philpott, Igor R. Efimov

## Abstract

Thermally-mediated ablation has utilized various energy sources, including cryothermal, radiofrequency (RF), microwave, laser, and high-frequency ultrasound with the goal of creating lesions to terminate focal sources or block reentrant wavefronts. RF- and cryo-ablation (CR) cause cell death through different mechanisms, and leave behind tissue with altered thermal-electric properties. We aimed to assess the effect of sequential RF and CR combinations on lesion size. Left ventricular (LV) wedge preparations (n=17) were dissected from ten donated human hearts and four epicardial ablation protocols were compared: 1) RF-RF (n=7); 2) CR-CR (n=7); 3) RF-CR (n=7); and 4) CR-RF (n=7). Preparations were continuously paced and perfused with oxygenated Tyrode solution. Ablated tissue was perfused for 3 hours, sectioned, and stained with 2,3,5-triphenyltetrazolium chloride to delineate necrosis. The effect of initial thermal-electric tissue properties on lesion depth during RF application was determined using a finite element method (FEM). CR-RF generated the deepest lesion (p<0.05) compared to protocols 1-3, while lesion width and area were similar among protocols. No energy combination produced a transmural lesion (n=0 of 28) in LV preparations. FEM showed that electrical conductivity plays a more significant role in lesion creation compared to thermal conductivity. A 33% increase in the initial thermal and electrical tissue conductivity generated a 21% deeper lesion. We conclude that sequential application of CR followed by RF created the deepest lesion in beating human LV preparations. This increase in lesion depth may translate into improved therapeutic outcomes for arrhythmias with intramural origins.

## Introduction

Ablation is an important therapeutic option to mitigate recurrent ventricular tachycardia (VT) that otherwise can give rise to ventricular fibrillation and sudden cardiac death.^1-6^ The goal of ablation is to terminate focal sources or block reentrant wavefronts that manifest in nonstructural and structural heart disease. Electrical mapping of focal origins allows for highly effective ablation targeting and usually requires small lesions to abolish the VT.^1, 6^ In structural heart disease, however, a surviving muscle isthmus hidden within the scar provides a critical segment for the reentrant circuit that can be difficult to consistently transect using ablation therapy.^1, 3, 7-9^ Intramural VT origin is a primary cause of ablation failure, and thus there is a need to increase lesion depth.^3, 10^

Cryothermal and radiofrequency (RF) ablation are the two most commonly used thermal modalities for treating arrhythmias. Both are highly effective at causing cellular death, but do so through different mechanisms. When temperature abruptly drops below −40°C during cryoablation (CR), intracellular water freezes causing irreversible disruption of membranes and intracellular organelles. During the early tissue re-warming phase, ice crystals coalesce into larger crystals further damaging membranes and organelles.^11-14^ Application of RF results in resistive heating of cardiac tissue proximal to the electrode, and thermal conduction contributes to the spread of tissue heating beyond the electrode-tissue interface. An increase in tissue temperature beyond +50°C causes permanent injury due to denaturing of proteins and coagulative necrosis.^15-17^

These mechanisms suggest that CR and RF ablation alter the electrical and thermal conductivity of tissue differently, which may be exploited to increase lesion size with a second application of energy. An increase in lesion depth would enhance the likelihood of transecting the critical segment of the reentrant circuit and improve ablation therapy of VT. In this study, we report lesion depth, width, and size after epicardial application of different CR-RF combinations in perfused human ventricular preparations. A finite element method (FEM) was used to further investigate the effect of initial thermal-electric tissue properties on lesion depth during RF application.

## Methods

### Donor Group

The study was approved by Institutional Review Board of Washington University School of Medicine (IRB ID 201105326). Human subjects gave informed consent. Donated hearts (n=10) came from two sources, Barnes-Jewish Hospital (Washington University School of Medicine) and Mid-America Transplant Services (Saint Louis, MO). Explanted hearts were arrested with cold (+4-7°C) cardioplegic solution (110 NaCl, 1.2 CaCl_2_, 16 KCl, 16 MgCl_2_, 10 NaHCO_3_ mmol/L) in the operating room.

### Experimental Preparations

Left ventricular (LV) wedge (n=17) preparations were dissected from donated human hearts as previously described.^18, 19^ Wedge preparations were isolated from several aspects of the LV free wall as permitted by available coronary arteries. Epicardial fat was carefully removed (Supplemental Figure I) to ensure direct contact between the energy source and the myocardium.

Preparations were perfused via left coronary arteries and endocardially submerged with warm (37°C), oxygenated (95% O_2_, 5% CO_2_) Tyrode’s solution (128.2 NaCl, 1.3 CaCl_2_, 4.7 KCl, 1.05 MgCl_2_, 1.19 NaH_2_PO_4_, 20 NaHCO_3_, 11.1 D-Glucose mmol/L). Tissue was perfused under constant pressure (60-80 mmHg) and perfusate pH was continuously monitored (7.35±0.05, Oakton Instruments, Vernon Hills, IL). Both the coronary perfusion and superfusion temperature were maintained at 37°C using a heating bath circulator (NESLAB EX7, Thermo Scientific, Asheville, NC). LV wedges were paced (PowerLab 26T, AD Instruments, Colorado Springs, CO) at 1 Hz at twice the diastolic threshold and Ag/AgCl pellet electrodes (WPI, Sarasota, FL) monitored (PowerLab 26T) the far-field ECG throughout experimentation. Tissue was allowed to stabilize for 20 min before initiation of an ablation protocol.

### Ablation Protocol

Ablation devices were positioned on the epicardial surface, while endocardial and transverse portions of the wedges were superfused with 37°C solution. Four epicardial ablation protocols were compared: 1) RF-RF (n=7); 2) CR-CR (n=7); 3) RF-CR (n=7); and 4) CR-RF (n=7). RF ablation was carried out using a Coolrail linear pen (AtriCure Inc., West Chester, OH) for 40 sec and CR was carried out with a cryoICE probe (AtriCure Inc., West Chester, OH) for 120 sec. Both probes were applied parallel to the tissue and there was a 2 min rest between applications. If wedge preparations were large enough for two lesions, the first ablation was applied distally and the second proximally to the cannula.

The Coolrail linear pen consists of two 30 mm long electrodes that are internally cooled, but not irrigated (Supplemental Figure IIA). Power is maintained at 30 W and is achieved within 2 sec. The cryoICE probe is 10 cm long with a 4.24 mm outer diameter (Supplemental Figure IIB); it is malleable, smooth, and has a thermal conductivity coefficient of 222 W/m.K. The probe is cooled by nitrous oxide (Joule-Thomson Principle) to −60°C and has an active defrost feature controlled by AtriCure Cryo Module (AtriCure, Inc., West Chester, OH).

### Tetrazolium Staining

Tissue was perfused for an additional 3 hours after completion of an ablation protocol to allow adequate time for lesion development. LV wedges were sectioned at ~2 mm intervals perpendicular to the ablation lesion creating transmural sections of myocardium. Sections from the periphery of the wedge were discarded, while 3-5 sections per wedge remained for staining. Sections were stained with 2,3,5-triphenyltetrazolium chloride (Sigma Aldrich, Saint Louis, MO) as described previously to delineate necrosis.^20^ Tetrazolium was diluted in a 2-part phosphate buffer comprising of NaHPO_4_ (0.10 mol/L) and NaH_2_PO_4_ (0.10 mol/L), which were combined in a ratio of 77.4%/22.6% (NaHPO_4_/NaH_2_PO_4_). Tetrazolium was added to the phosphate buffer at 1% weight/volume (gm/mL). All sections were incubated for 15-20 min at 37°C and then photographed.

### Lesion Quantification

Images of tetrazolium-stained sections were converted to an 8-bit scale and thresholded to delineate lesions. Lesion depth, width, and surface area were quantified for all sections within a wedge and then averaged using custom-built software (Matlab 2012b, Mathworks).

### Computational RF Ablation Model

Pre-ablated tissue has different thermo-electric conductivities than post-ablated tissue. We used FEM (COMSOL Multiphysics, Palo Alto, CA) to develop a computer model to test the effect of the initial thermal and electrical tissue conductivities on lesion creation during RF ablation. The model was designed to mimic the epicardial ablation that was utilized in the in-vitro experimental conditions.

#### Governing Equations

The conversion of electric energy into heat injures tissue. Joule heating will cause the temperature of the myocardium near the ablation electrode to rise and thermal conduction will cause the temperature to increase deeper within the tissue. The temperature distribution in the modeled myocardium was determined by solving the bio-heat equation:^21^

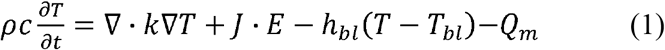

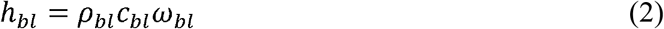

where T is temperature distribution, ρis density (kg/m^3^), c is heat capacity (J/kg.K), k is thermal conductivity (W/m.K), J is current density (A/m^2^), and E is the electric field intensity (V/m). The subscript *bl* differentiates blood properties from myocardium, where h_bl_ (W/m^2^.K) is the convective heat transfer coefficient of blood. The perfusion of blood within the ventricular cavity, which acts as a heat sink, is characterized by the term ω (1/s). The metabolic energy Q_m_ generated by the beating heart is set to 0 W/m^3^.^22^

Equation (1) is solved in a few steps. First, the potential distribution is determined by solving Laplace equation (3).

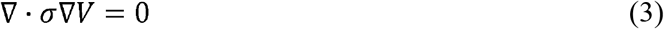

where V is the potential distribution (V) and σis electrical conductivity (S/m). A 500 kHz, 30 V RF ablation was applied at one electrode, while the other served as ground. After V is calculated, E is computed from (4) and J from (5)

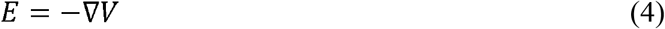

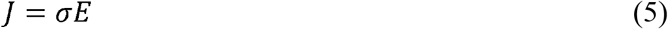

Next, the temperature distribution T is solved using Equation (1). Lesion size was determined from the +50°C contour after 40 sec of RF application.

#### Geometry and Boundary Conditions

The geometry of the heart was simplified to a two-dimensional region, 15 x 20 mm, or the average transverse section of the LV wedge preparation (Figure 1A) we created. The RF ablation electrodes penetrate the myocardium 0.40 mm and have a 4 mm gap between them. There is a thermal insulation layer between electrodes to resemble their plastic encapsulation. The endocardial surface is in contact with circulating blood (37°C) and acts as a heat sink.

**Figure 1:**
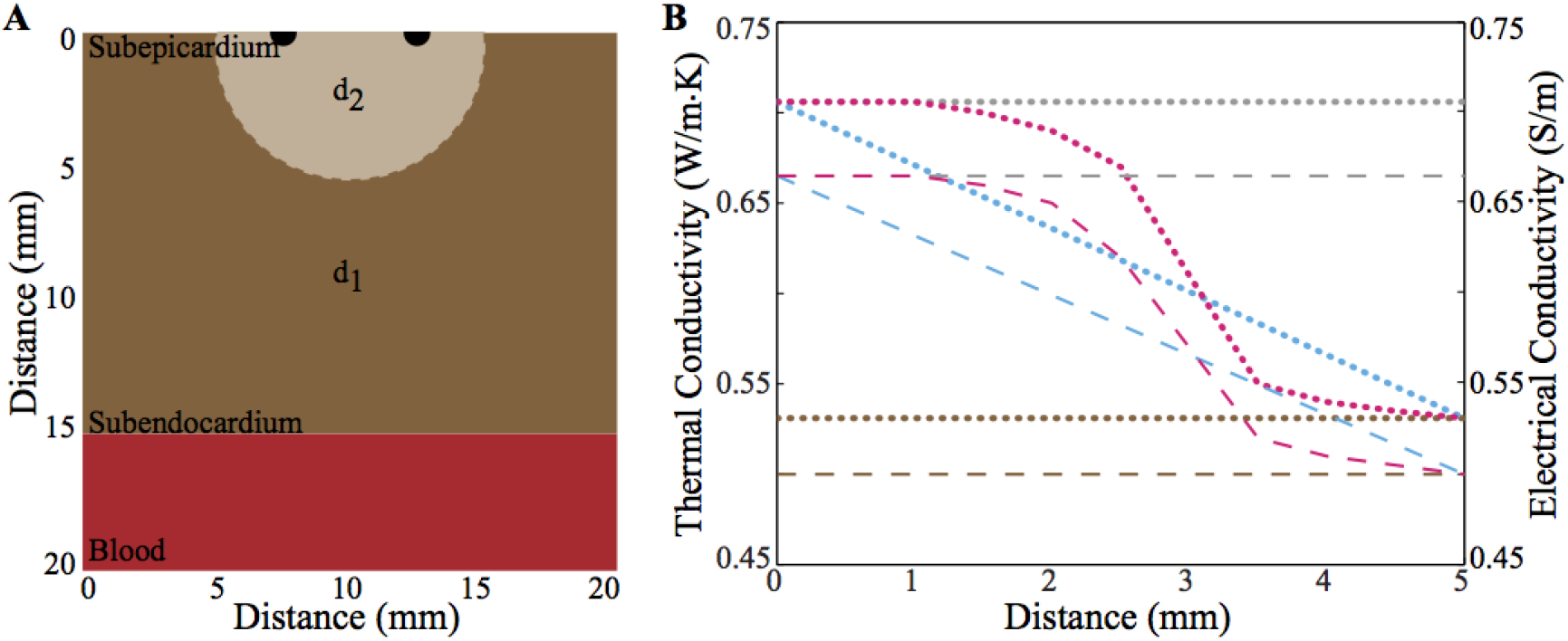
Model geometry and spatially dependent conductivities for radiofrequency ablation. **(A)** Model geometry consists of myocardium (brown), electrodes (black), and blood (red). The myocardium has two domains d_1_ and d_2_, whose electrical (σ) and thermal (κ) conductivities are plotted in panel B. **(B)** d_1_ values are plotted brown, while d_2_ values are plotted brown (d_2_ = d_1_), grey, teal, and pink. Dashed lines are σ values and dotted lines k values.

The following are the initial and boundary conditions:

> T=37°C when t=0 sec in entire domain
>
> T=37°C when t≥0 sec at transverse boundaries

The parameter values for myocardium, blood, and the electrodes are presented in Supplementary Table I.^23, 24^ The temperature-dependence of tissue properties was ignored during the 40 sec ablation.

The myocardium is separated into two domains, d_1_ and d_2_. Domain d_1_ is far away from the ablation electrode and has constant thermal conductivity of 0.531 W/m.K and electrical conductivity of 0.50 S/m, while d_2_ is underneath the ablation electrodes and has varying tissue properties (Figure 1B). Domain d_2_ is a semicircle with a depth of 5 mm and width of 10 mm. The temperature within the myocardium was determined when the thermal and electrical conductivities of d_2_ were: d_2_ = d_1_ (brown), d_2_ = 1.33.d_1_ (grey), a linear decrease from 1.33.d_1_ to d_1_ (teal), and a sigmoidal decay from 1.33.d_1_ to d_1_ (pink). When electrical and thermal conductivities were tested separately within d_2_, the other parameter was set to the corresponding d_1_ value.

### Statistics

All data are expressed as the mean ± standard error of the mean. A p-value of 0.05 was considered significant. Statistically significant differences were identified using a oneway analysis of variance (ANOVA). A post-hoc Tukey test was performed if ANOVA was significant.

## Results

Figure 2A shows a representative LV wedge preparation that was ablated (Left), stained (Middle), and segmented (Right). The top lesion (teal) is from a CR-RF ablation, while the bottom lesion (pink) is from a RF-CR ablation. Tetrazolium staining clearly identifies necrotic tissue (white) from viable tissue (red), which is segmented for analysis. Lesion depth, width, and area were quantified for all ablation protocols (Figure 2B-D). CR-RF generated a significantly greater (p<0.05) lesion depth of 7.60±0.31 mm, while RF-RF, CR-CR, and RF-CR created depths of 5.38±0.37 mm, 6.23±0.57 mm, and 6.24±0.33 mm, respectively. Lesion width did not differ among protocols (p=0.67). CR-RF additionally generated the largest lesion area of 80.52±6.60 mm^2^, while RF-RF, CR-CR, and RF-CR created lesion areas of 52.67±1.75 mm^2^, 69.34±9.14 mm^2^, and RF-CR 57.21±7.67 mm^2^, respectively. ANOVA analysis of lesion area produced a p-value of 0.043, but a post-hoc Tukey test determined lesion areas were not different.

**Figure 2:**
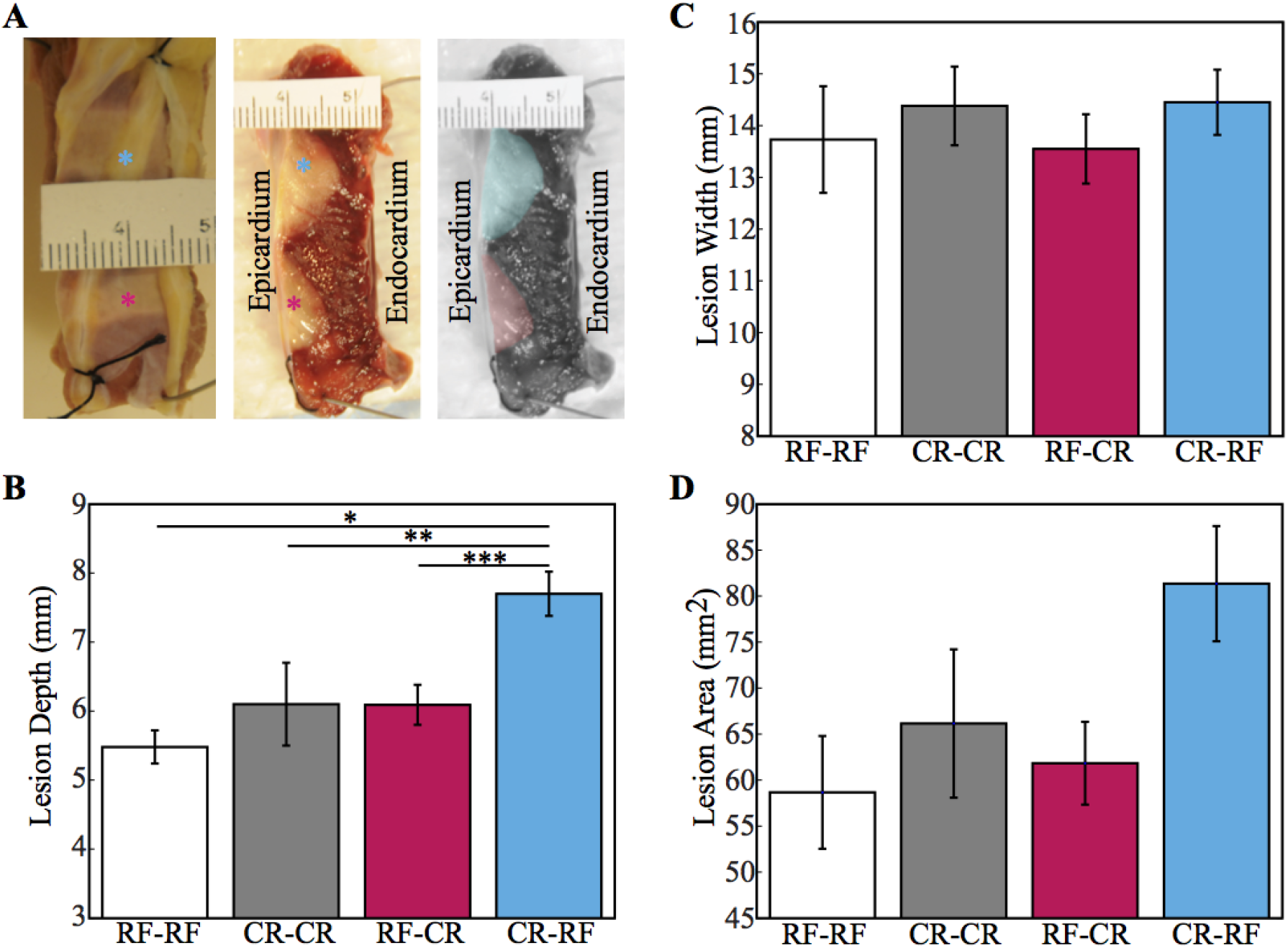
Lesion quantification during different combinations of cryoablation (CR) and radiofrequency (RF) ablation. **(A)** Ablated preparation (Left), stained transverse section of tissue (Middle), and digitally segmented lesions (Right). CR-RF created top lesion (teal) and RF-CR created bottom lesion (pink). **(B)** Lesion depth. **(C)** Lesion width **(D)** Lesion area. *CR-RF versus RF-RF p<0.05, **CR-RF versus CR-CR p<0.05, ***CR-RF versus RFCR p<0.05

LV wedge preparations were 14.9±0.6 mm thick and ranged from 9.9 mm to 21.8 mm. Figure 3 displays a scatter plot of tissue thickness versus lesion depth for all n=28 ablations. No energy combination produced a transmural lesion (n=0 of 28), as highlighted by no circles being on the dotted line. The scatter plot also indicates that tissue thickness did not influence lesion depth. Lesion creation in thin ventricular tissue (~10 mm) was not impacted by the endocardial 37°C heat sink compared to thicker tissue. The R-squared values are 0.01 for RF-RF, 0.002 for CR-CR, 0.49 for RF-CR, and 0.03 for CR-RF.

**Figure 3:**
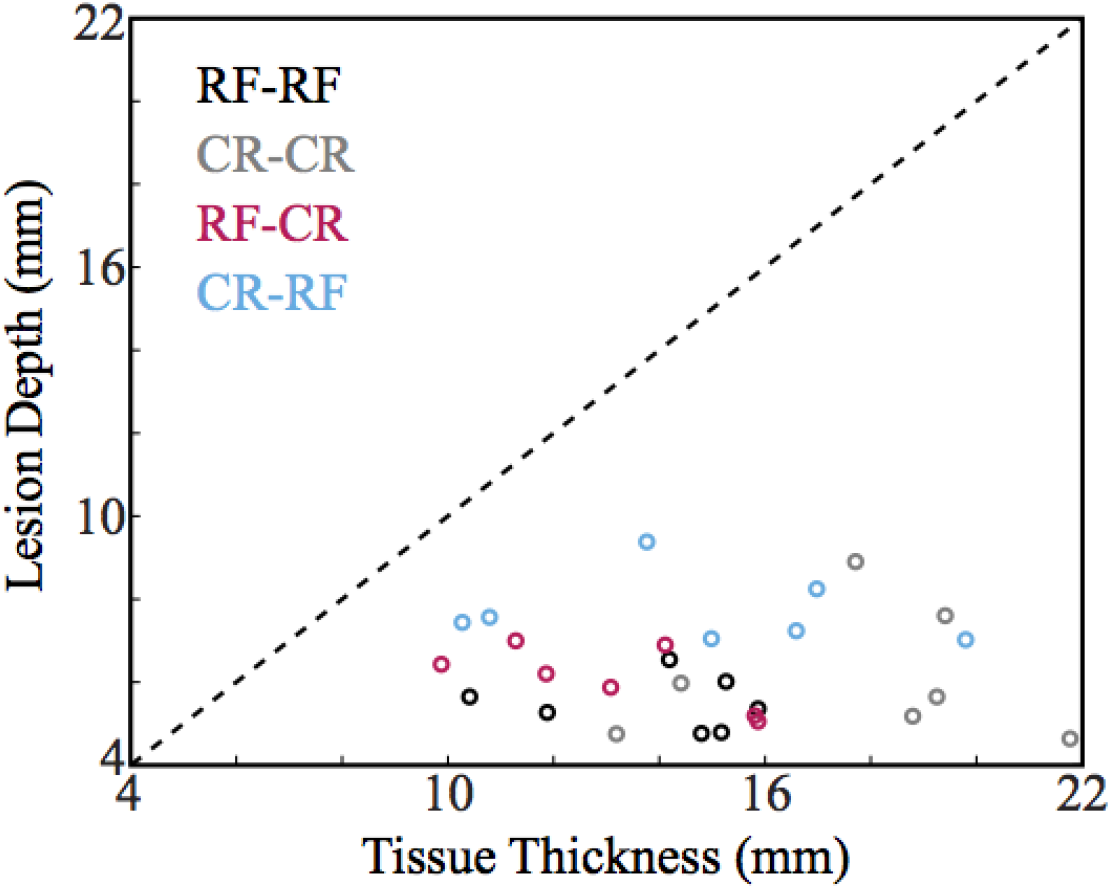
Lesion depth in relation to tissue thickness.CR= Cryoablation, RF= Radiofrequency Ablation

The initial electrical and thermal conductivities of cardiac tissue play a critical role in determining lesion extent when applying RF energy. An FEM analysis was employed with spatially varying thermal-electric tissue properties (Figure 1B) to quantify their effect on lesion size. Figure 4A displays the temperature distribution after 5, 10, 20, and 40 sec of RF application when d_2_ conductivities decay sigmoidally from the epicardial surface. A large temperature increase is seen proximal to the ablation electrodes, which spreads deeper into the tissue. At 40 sec thermal conduction is unable to increase the initial temperature of the subendocardium above 37°C.

**Figure 4:**
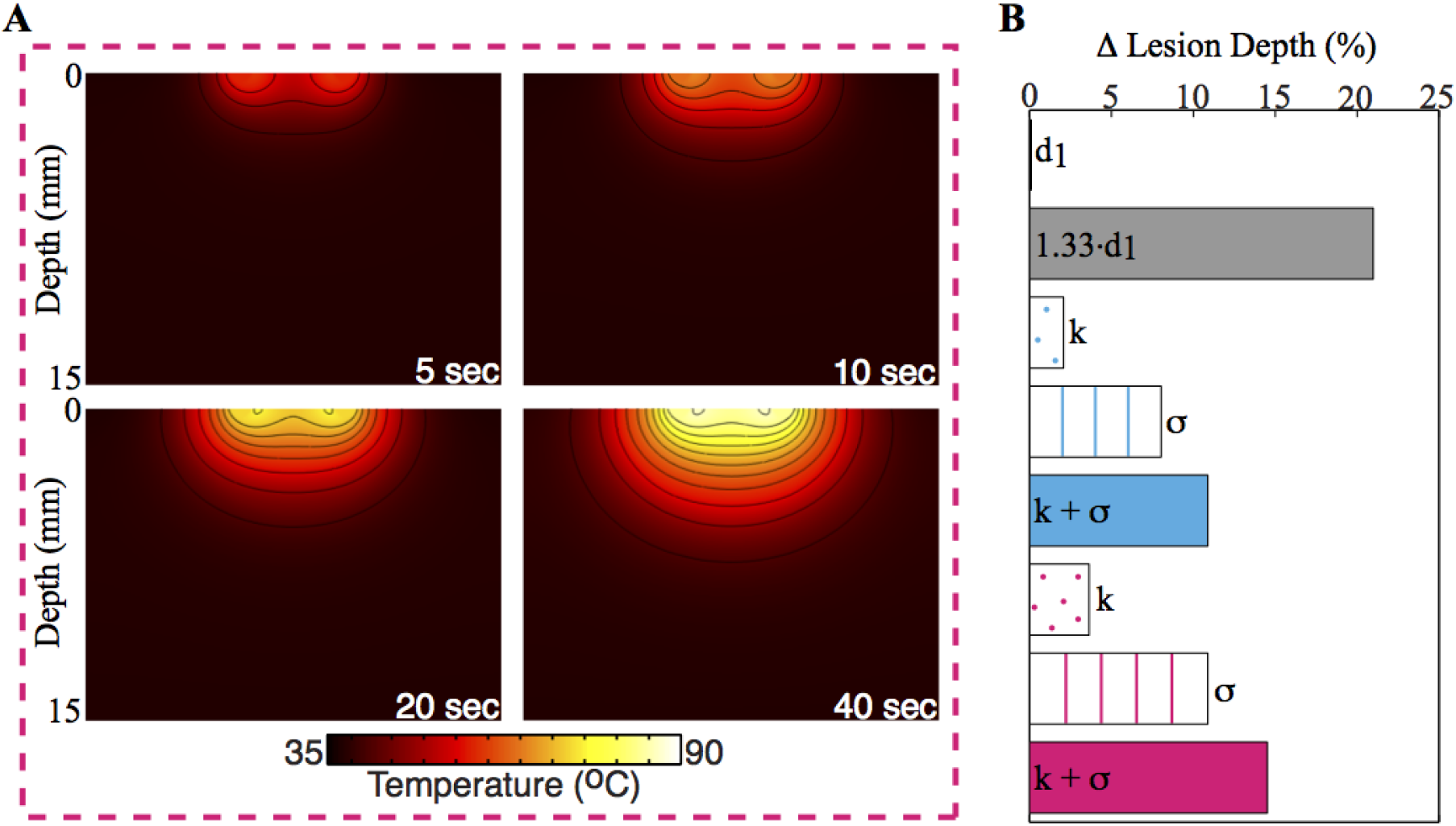
Modeled temperature distribution and lesion depth. **(A)** Example temperature profile at 5, 10, 20, and 40 sec using sigmoidal d_2_ properties (pink). Temperature isochrones are from 40°C to 90°C with steps of 5°C. **(B)** Percent change in lesion depth when using d_2_ properties. d_1_=Domain 1, d_2_=Domain 2, k=Thermal Conductivity, σ = Electrical Conductivity

When d_2_=1.33.d_1_ (grey), lesion depth increased by 21.0% compared to d_2_=d_1_, while the linear (teal) and sigmoidal (pink) changes in conductivities increased the lesion depth by 10.9% and 14.5%, respectively (Figure 4B). Electrical conductivity played a more significant role than thermal conductivity in lesion creation. When thermal conductivity was held at its d_1_ value (0.531 W/m.K) and the electrical conductivity changed linearly and sigmoidally, lesion depth increased 8.0% (σ-blue lined) and 10.9% (σ-pink lined) respectively; however, when electrical conductivity was held at its d_1_ value (0.50 S/m) and thermal conductivity changed linearly and sigmoidally, lesion depth increased only by 2.1% (k-blue dotted) and 3.6% (k-pink dotted). This demonstrates that lesion depth is primarily controlled by electrical conductivity.

## Discussion

Several studies have investigated using multiple lesion lines to improve ablation efficacy or their necessity to close lesion gaps using the same energy source.^25-28^ This is the first study to examine of the effect of multiple energy sources on lesion size in the human heart. Implementation of CR and then RF generates a significantly larger lesion depth compared to all other RF-CR combinations. The lesion width created by all RF-CR combinations was similar as well as the total lesion area. The substantial thickness of the human LV prevented any energy combinations from generating transmural lesions.

A hybrid transcatheter system has previously been developed and tested in an in-vivo canine model that is capable of delivering RF and CR independently, sequentially, and simultaneously.^29^ Khairy et al elegantly showed that simultaneous application of CR (10°C) and RF (45 W) produced a greater lesion depth (6.7 mm) compared to both standard RF ablation (5.5 mm) and irrigated RF ablation (5.4 mm). Dissimilar to our findings, the sequential application of CR (−75°C) followed by RF (20 W) created a smaller lesion depth (4.6 mm). A possible explanation for the difference could be due to our utilization of a 2 min thawing period prior to RF application. In the early thawing phase, intracellular ice crystals coalescence and increase in size. This further disrupts cellular membranes decreasing tissue impedance for the subsequent RF to penetrate deeper into the tissue. Application of RF while the tissue is frozen may limit the utility of CR-RF.

To our knowledge cardiac tissue properties after CR and RF are unknown. Previously Haemmerich and colleagues^30^ measured the tissue properties after RF on a single excised human liver sample and showed a sizeable 270% increase in tissue conductivity at 10 Hz, but only a modest 10% increase in tissue conductivity near the RF range (1 MHz). We utilized an FEM simulation to test the effect of the initial tissue properties on RF application. Tungjitkusolmun et al^22, 31^ previously did innovative RF modeling and simulated the effect of changing myocardial tissue properties by 50 and 100%;^24^ however, neither the spatial dependency of the properties nor the epicardial application of RF in ventricular tissue has been examined. We chose a maximum increase of 33% to keep conductivities in a physiological range for cardiac tissue. We demonstrated a lesion depth increase of 21%, which is primarily driven by the initial electrical conductivity.

Experimentally, CR-RF application enhanced lesion depth by 40.5% compared to RF-RF, and possibly other factors besides tissue properties contribute to CR-RF’s effectiveness. Ice crystal formation during CR is known to shear organelles, including the sarcoplasmic reticulum. Upon re-warming, the increased cytosolic Ca^2+^ concentration may result in hypercontracture, or maintained sarcomere shortening. Hypercontracture would increase the number of myocytes in a given volume of tissue for the subsequent RF application, which would enhance the amount of tissue injured. Alternatively, thawing tissue temperatures may act similarly to an irrigated-tip RF ablation and enhance lesion depth.^32^

### Limitations

We acknowledge a few important limitations to this study. First, all RF-CR protocols were performed on contracting ex vivo wedge preparations that were perfused by Tyrode’s solution. The experimental setup, although similar, does not entirely mimic the tension and motion of an intact beating human heart. In addition, we did not control for contact force. Previous studies^33, 34^ have shown that increases in contact force will increase lesion size. Haines et al reported that contact force becomes insignificant for forces greater than 0.10 N as long as an adequate tissue-catheter interface temperature is maintained.^35^ During epicardial ablation a contact force of 0.10 N is easily achieved. In studies where contact force is directly related to lesion size, an increase in force is associated with both an increase in lesion depth and width. In this study all RF-CR combinations had similar lesion widths and therefore the differences in lesion depths are most likely not related to dissimilar contact forces.

### Clinical Implication

Ablation is an essential therapeutic option to decrease recurrent ventricular tachycardia that otherwise can lead to ventricular fibrillation and sudden cardiac death. Intramural ventricular tachycardia origins require lesions to penetrate deep within the ventricular tissue. Cryoablation followed by radiofrequency ablation generated the deepest lesion compared to all other combinations and may be more adept at transecting the surviving muscle isthmus hidden within the scar.

## Conclusion

This is the first study to investigate lesion size using multiple energy sources in the human heart. Cryoablation followed by radiofrequency ablation generated the largest lesion compared to all other combinations in beating, perfused human LV preparations. Cryo-radiofrequency ablation could have important implications for surgical treatment of structural heart disease, where the surviving muscle isthmus in the ventricles can be difficult to transect. The exact mechanism for the increase in lesion depth still needs to be elucidated; likely, cryoablation increases the thermal-electric properties of the myocardium and thus enhances the efficacy of radiofrequency ablation.

## Acknowledgement

NIH grants R01-HL115415 and R01-HL114395 and grant RHYTHM from the Leducq Foundation are gratefully acknowledged.

## List of Abbreviations

CR—: Cryoablation
FEM—: Finite Element Method
LV—: Left Ventricle
RF—: Radiofrequency
VT—: Ventricular Tachycardia

## References

1. Aliot EM, Stevenson WG, Almendral-Garrote JM, et al. EHRA/HRS Expert Consensus on Catheter Ablation of Ventricular Arrhythmias: developed in a partnership with the European Heart Rhythm Association (EHRA), a Registered Branch of the European Society of Cardiology (ESC), and the Heart Rhythm Society (HRS); in collaboration with the American College of Cardiology (ACC) and the American Heart Association (AHA). Europace : European pacing, arrhythmias, and cardiac electrophysiology : journal of the working groups on cardiac pacing, arrhythmias, and cardiac cellular electrophysiology of the European Society of Cardiology Jun 2009;11:771-817.

2. Klein LS, Shih HT, Hackett FK, Zipes DP, Miles WM. Radiofrequency catheter ablation of ventricular tachycardia in patients without structural heart disease. Circulation May 1992;85:1666-1674.

3. Tokuda M, Kojodjojo P, Tung S, Tedrow UB, Nof E, Inada K, Koplan BA, Michaud GF, John RM, Epstein LM, Stevenson WG. Acute failure of catheter ablation for ventricular tachycardia due to structural heart disease: causes and significance. Journal of the American Heart Association Jun 2013;2:e000072.

4. Coggins DL, Lee RJ, Sweeney J, Chein WW, Van Hare G, Epstein L, Gonzalez R, Griffin JC, Lesh MD, Scheinman MM. Radiofrequency catheter ablation as a cure for idiopathic tachycardia of both left and right ventricular origin. Journal of the American College of Cardiology May 1994;23:1333-1341.

5. Calkins H, Epstein A, Packer D, et al. Catheter ablation of ventricular tachycardia in patients with structural heart disease using cooled radiofrequency energy: results of a prospective multicenter study. Cooled RF Multi Center Investigators Group. Journal of the American College of Cardiology Jun 2000;35:1905-1914.

6. Haissaguerre M, Shoda M, Jais P, et al. Mapping and ablation of idiopathic ventricular fibrillation. Circulation Aug 20 2002;106:962-967.

7. Stevenson WG, Wilber DJ, Natale A, et al. Irrigated radiofrequency catheter ablation guided by electroanatomic mapping for recurrent ventricular tachycardia after myocardial infarction: the multicenter thermocool ventricular tachycardia ablation trial. Circulation Dec 16 2008;118:2773-2782.

8. Carbucicchio C, Santamaria M, Trevisi N, Maccabelli G, Giraldi F, Fassini G, Riva S, Moltrasio M, Cireddu M, Veglia F, Della Bella P. Catheter ablation for the treatment of electrical storm in patients with implantable cardioverter-defibrillators: short- and long-term outcomes in a prospective single-center study. Circulation Jan 29 2008;117:462-469.

9. Tanner H, Hindricks G, Volkmer M, Furniss S, Kuhlkamp V, Lacroix D, C DEC, Almendral J, Caponi D, Kuck KH, Kottkamp H. Catheter ablation of recurrent scar-related ventricular tachycardia using electroanatomical mapping and irrigated ablation technology: results of the prospective multicenter Euro-VT-study. Journal of cardiovascular electrophysiology Jan 2010;21:47-53.

10. Gizurarson S, Spears D, Sivagangabalan G, Farid T, Ha AC, Masse S, Kusha M, Chauhan VS, Nair K, Harris L, Downar E, Nanthakumar K. Bipolar ablation for deep intra-myocardial circuits: human ex vivo development and in vivo experience. Europace : European pacing, arrhythmias, and cardiac electrophysiology : journal of the working groups on cardiac pacing, arrhythmias, and cardiac cellular electrophysiology of the European Society of Cardiology Feb 19 2014.

11. Gage AA, Baust J. Mechanisms of tissue injury in cryosurgery. Cryobiology Nov 1998;37:171-186.

12. Skanes AC, Yee R, Krahn AD, Klein GJ. Cryoablation of atrial arrhythmias. Cardiac electrophysiology review Dec 2002;6:383-388.

13. Cooper IS. CRYOBIOLOGY AS VIEWED BY THE SURGEON. Cryobiology Sep-Oct 1964;51:44-51.

14. Baust JG, Gage AA, Bjerklund Johansen TE, Baust JM. Mechanisms of cryoablation: clinical consequences on malignant tumors. Cryobiology Feb 2014;68:1-11.

15. Kalbfleisch SJ, Langberg JJ. Catheter Ablation with Radiofrequency Energy: Biophysical Aspects and Clinical Applications. Journal of cardiovascular electrophysiology 1992;3:173-186.

16. Avitall B, Khan M, Krum D, Hare J, Lessila C, Dhala A, Deshpande S, Jazayeri M, Sra J, Akhtar M. Physics and engineering of transcatheter cardiac tissue ablation. Journal of the American College of Cardiology Sep 1993;22:921-932.

17. Goldberg SN, Gazelle GS, Mueller PR. Thermal ablation therapy for focal malignancy: a unified approach to underlying principles, techniques, and diagnostic imaging guidance. AJR American journal of roentgenology Feb 2000;174:323-331.

18. Lou Q, Fedorov VV, Glukhov AV, Moazami N, Fast VG, Efimov IR. Transmural heterogeneity and remodeling of ventricular excitation-contraction coupling in human heart failure. Circulation May 3 2011;123:1881-1890.

19. Glukhov AV, Fedorov VV, Kalish PW, Ravikumar VK, Lou Q, Janks D, Schuessler RB, Moazami N, Efimov IR. Conduction remodeling in human end-stage nonischemic left ventricular cardiomyopathy. Circulation Apr 17 2012;125:1835-1847.

20. Ytrehus K, Liu Y, Tsuchida A, Miura T, Liu GS, Yang XM, Herbert D, Cohen MV, Downey JM. Rat and rabbit heart infarction: effects of anesthesia, perfusate, risk zone, and method of infarct sizing. The American journal of physiology Dec 1994;267:H2383-2390.

21. Pennes HH. Analysis of tissue and arterial blood temperatures in the resting human forearm. 1948. Journal of applied physiology (Bethesda, Md : 1985) Jul 1998;85:5-34.

22. Tungjitkusolmun S, Staelin ST, Haemmerich D, Tsai JZ, Webster JG, Lee FT, Jr., Mahvi DM, Vorperian VR. Three-Dimensional finite-element analyses for radio-frequency hepatic tumor ablation. IEEE transactions on bio-medical engineering Jan 2002;49:3-9.

23. Panescu D, Whayne JG, Fleischman SD, Mirotznik MS, Swanson DK, Webster JG. Three-dimensional finite element analysis of current density and temperature distributions during radio-frequency ablation. IEEE transactions on bio-medical engineering Sep 1995;42:879-890.

24. Tungjitkusolmun S, Woo EJ, Cao H, Tsai JZ, Vorperian VR, Webster JG. Thermal–electrical finite element modelling for radio frequency cardiac ablation: effects of changes in myocardial properties. Medical & biological engineering & computing Sep 2000;38:562-568.

25. Schneider HE, Stahl M, Kriebel T, Schillinger W, Schill M, Jakobi J, Paul T. Double cryoenergy application (freeze-thaw-freeze) at growing myocardium: lesion volume and effects on coronary arteries early after energy application. Implications for efficacy and safety in pediatric patients. Journal of cardiovascular electrophysiology Jun 2013;24:701-707.

26. Laughner JI, Sulkin MS, Wu Z, Deng CX, Efimov IR. Three potential mechanisms for failure of high intensity focused ultrasound ablation in cardiac tissue. Circulation Arrhythmia and electrophysiology Apr 2012;5:409-416.

27. Soejima K, Suzuki M, Maisel WH, Brunckhorst CB, Delacretaz E, Blier L, Tung S, Khan H, Stevenson WG. Catheter ablation in patients with multiple and unstable ventricular tachycardias after myocardial infarction: short ablation lines guided by reentry circuit isthmuses and sinus rhythm mapping. Circulation Aug 7 2001;104:664-669.

28. Marchlinski FE, Callans DJ, Gottlieb CD, Zado E. Linear ablation lesions for control of unmappable ventricular tachycardia in patients with ischemic and nonischemic cardiomyopathy. Circulation Mar 21 2000;101:1288-1296.

29. Khairy P, Cartier C, Chauvet P, Tanguay JF, Simeon B, Lalonde JP, Dubuc M. A novel hybrid transcatheter ablation system that combines radiofrequency and cryoenergy. Journal of cardiovascular electrophysiology Feb 2008;19:188-193.

30. Haemmerich D, Schutt DJ, Wright AW, Webster JG, Mahvi DM. Electrical conductivity measurement of excised human metastatic liver tumours before and after thermal ablation. Physiological measurement May 2009;30:459-466.

31. Tungjitkusolmun S, Vorperian VR, Bhavaraju N, Cao H, Tsai JZ, Webster JG. Guidelines for predicting lesion size at common endocardial locations during radio-frequency ablation. IEEE transactions on bio-medical engineering Feb 2001;48:194-201.

32. Nakagawa H, Yamanashi WS, Pitha JV, Arruda M, Wang X, Ohtomo K, Beckman KJ, McClelland JH, Lazzara R, Jackman WM. Comparison of in vivo tissue temperature profile and lesion geometry for radiofrequency ablation with a saline-irrigated electrode versus temperature control in a canine thigh muscle preparation. Circulation Apr 15 1995;91:2264-2273.

33. Wong MC, Edwards G, Spence SJ, Kalman JM, Kumar S, Joseph SA, Morton JB. Characterization of catheter-tissue contact force during epicardial radiofrequency ablation in an ovine model. Circulation Arrhythmia and electrophysiology Dec 2013;6:1222-1228.

34. Yokoyama K, Nakagawa H, Shah DC, Lambert H, Leo G, Aeby N, Ikeda A, Pitha JV, Sharma T, Lazzara R, Jackman WM. Novel contact force sensor incorporated in irrigated radiofrequency ablation catheter predicts lesion size and incidence of steam pop and thrombus. Circulation Arrhythmia and electrophysiology Dec 2008;1:354-362.

35. Haines DE. Determinants of Lesion Size During Radiofrequency Catheter Ablation: The Role of Electrode-Tissue Contact Pressure and Duration of Energy Delivery. Journal of cardiovascular electrophysiology 1991;2:509-515.

